# Ecophylogenetics Reveals the Evolutionary Associations between Mammals and their Gut Microbiota

**DOI:** 10.1101/182212

**Authors:** Christopher A. Gaulke, Holly K. Arnold, Steven W. Kembel, James P. O’Dwyer, Thomas J. Sharpton

**Author notes:** Corresponding Author: Thomas J. Sharpton.

## Abstract

A tantalizing hypothesis posits that mammals coevolved with their gut microbiota. Unfortunately, the limited resolution of microbial taxonomy hinders the exploration of this hypothesis and specifically challenges the discovery of gut microbes that are linked to mammalian evolution. To address this, we developed a novel approach that groups microbes into new, more meaningful taxonomic units based on their common ancestry and ecological redundancy. Treating mammalian lineages as different ecosystems, we quantified the distribution of these microbial taxa across mammals. Our analysis discovered monophyletic clades of gut bacteria that are unexpectedly prevalent, or conserved, across all mammals, as well as conserved clades that are exclusive to particular mammalian lineages. These clades often manifest phylogenetic patterns indicating that they are subject to selection. Lineage - specific changes in clade conservation, including a human-accelerated loss of conserved clades, suggest that mammalian evolution associates with a change in the selective regimes that act on gut microbiota. Collectively, these results point to the existence of microbes that possess traits that facilitate their dispersion or survival in the mammalian gut, possibly because they are subject to host selection. Ultimately, our analysis clarifies the relationship between the diversification of the gut microbiome and mammalian evolutionary history.

## Introduction

Emerging research links the gut microbiome to mammalian evolution, but there remains limited insight into the processes underlying this association. Trillions of microbes inhabit the mammalian gut and perform vital functions for their host that could modulate niche specificity or survival, such as nutrient acquisition, biosynthesis and detoxification (1, 2), stimulation of gut and immune development (3), and defense against infection (4). The gut microbiome may also affect ecophysiologically relevant aspects of behavior and stress response (5). Consequently, gut microbiota may influence host fitness and natural selection may favor hosts that harbor specific assemblages of gut microbiota. Furthermore, mammalian evolution yielded changes in physiology, behavior, diet, or ecological niche that could influence which microbes are exposed to the gut (i.e., the metacommunity) or that can thrive within it. The microbiome could also be vertically inherited, either directly or as a result of host genotypic filtering of environmentally acquired microbes. In support of these possibilities, recent studies have found that the biodiversity of the gut microbiome correlates with mammalian phylogeny (i.e., phylosymbiosis) (6 – 8). However, there is debate about the contribution of potential confounding factors. Differences in diet (8, 9), intestinal morphology, management facility (10), or the local metacommunity (11) could influence the observed variation across lineages. That said, studies have identified a limited number of mammalian gut microbiota that have co - diversified with their hosts (8, 12), are subject to vertical inheritance (13, 14), or that associate with host genotype (14, 15). These findings indicate that host genomic evolution associates with at least some of the variation in the gut microbiome.

Quantifying the distribution of enteric microbes among mammals illuminates the processes underlying this association. For example, microbiota that are common to mammals may be keystone members of the microbiome (16), apt gut generalists, or critical to mammalian fitness and subject to selection. Additionally, microbiota that associate with specific mammalian taxonomies may be sensitive to properties derived in the mammalian ancestor, including changes in physiology, diet, behavior, or niche. These properties may impact neutral (e.g., metacommunity assembly) or selective (e.g., physiological filtering) processes that influence the microbe‘s presence in the gut. These properties may also influence how a mammal‘s fitness depends upon the biological functions executed by the microbe. Only by first determining how microbial taxa distribute among mammals can we discern the specific ecological and evolutionary processes that underlie their association.

Prior efforts to define this distribution are complicated by the diffuse nature of microbial taxonomy. While the classification of microbes into a Linnaean taxonomy (i.e., phylotyping) detects associations between broad taxonomic categories and ecological or host covariates, this approach cannot resolve differences in intermediate levels of taxonomy. Consequently, it fails to identify associations that are complicated by phylogenetic redundancy, wherein multiple phylotypes descended from a common ancestor and share synapomorphic traits that underlie their ecological distribution. As a result, these phylotypes are functionally interchangeable across communities and statistical tests that operate at the level of distinct phylotypes may fail to resolve associations due to problems of sparsity. Unfortunately, analyses at higher order levels of taxonomy do not necessarily solve this problem because of phylogenetic aggregation: higher order phylotypes will not only include the ancestor from which the traits in question derived, but also other lineages that do not possess the traits. Consequently, tests of association may fail to detect a signal amidst the noise. These challenges likely confound the resolution of gut microbes that are shared across mammals, given that 1) few enteric microbial Operational Taxonomic Units (OTUs) are shared across mammalian species, while higher order phylotypes are (7), and 2) phylogenetic relationships often predict microbial trait conservation (17 – 19).

To circumvent these challenges, we introduce a phylogenetically flexible taxonomy that groups microbial lineages into ecologically relevant taxonomic units (Figure 1). In particular, we adopt ecophylogenetic theory (20) to identify taxa as monophyletic clades that manifest statistical associations with ecological covariates. These covariates could include quantitative characteristics, such as environmental pH, or categorical characteristics, such as whether the community was sampled from a marine versus terrestrial environment. For example, a clade whose subtending lineages collectively stratify communities in association with a categorical characteristic would represent a taxonomic unit.

**Figure 1.**
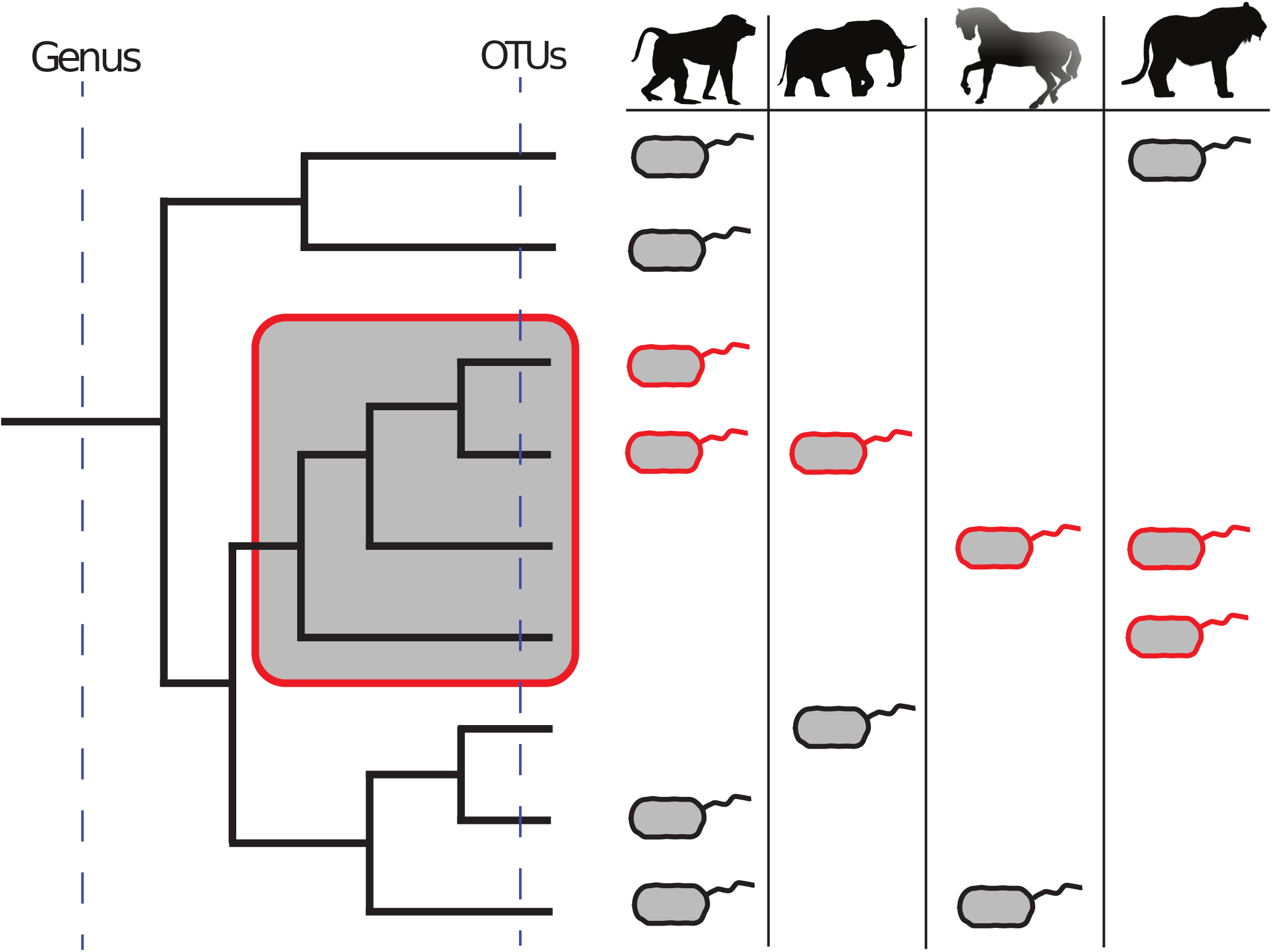
An ecophylogenetic approach to taxonomy can discover ecologically relevant units of microbial taxa. Incorporating phylogeny into the assessment of how microbial lineages are distributed across communities can identify monophyletic clades of microbes that collectively manifest an association with ecological factors. For example, the clade highlighted in red is universally present across all mammalian microbiome samples, indicating that the clade may have evolved a conserved trait that facilitated its ubiquitous distribution. If we were to consider this relationship at the OTU level (*i.e.,* considering the tips of the tree as appropriate units), the redundancy of OTUs within this clade would obscure the detection of this relationship. On the other hand, if we were to consider the genus level, the aggregation of this clade with others that do not possess the trait would similarly obscure this relationship.

Our taxonomic approach benefits from several features. First, it relies on phylogenetic relationships, so it is not biased by contemporary taxonomic labels and is not confounded by phylogenetic redundancy or aggregation. In addition, the phylogeny provides an opportunity to assess whether a clade‘s prevalence across samples is due to chance or not. For example, ancient clades are more likely to contain lineages in a diverse set of communities. Conversely, recently emerged clades may not be found in all communities, but in a greater number than expected by chance, indicating that a non - random process influenced their ecological diversification. Furthermore, clades that associate with ecological covariates represent the theoretical evolutionary origin of the microbial traits that underlie the ecological association. For example, clades that are common across mammals likely derived traits in their ancestor that are critical to the function of the microbial community, the fitness of the host, or the ability of the microbes to disperse and succeed within the host gastrointestinal tract. We note that this concept is theoretically consistent with the ecotype model of speciation, which posits that speciation results from adaptation to local ecological conditions (21).

We used this definition of taxonomy to clarify how gut microbes associate with mammalian evolutionary history. Specifically, we treated the mammalian gut (or the gut of specific mammalian lineages) as an ecosystem and applied our taxonomic approach to resolve microbial taxa that are more prevalent among mammals (or specific mammalian lineages) than expected by chance. We refer to these taxa as conserved clades given that non - random processes have resulted in their ecological prevalence being conserved across mammalian species. Our analysis reveals that conserved clades were integrated into or lost from mammalian guts in a manner correlated with mammalian evolutionary history. Furthermore, conserved clades manifest evolutionary patterns consistent with being subject to selection, possibly because they have coevolved with their hosts. These results demonstrate the value of our taxonomic approach and reveal how selective processes have influenced the diversification of the mammalian gut microbiome.

## Results

### Mammalian evolution associates with conserved clades of bacteria

We developed an algorithm and corresponding software (ClaaTU) that identifies conserved monophyletic clades of taxa, which are clades that are more prevalent across a defined set of communities than expected by chance. Briefly, our procedure traverses a phylogeny assembled from 16S sequences that were generated from multiple communities. It then quantifies each clade‘s prevalence across a defined subset of the communities, where the clade‘s prevalence is based on the occurrence of the subtending lineages in the subset of communities. A permutation test then quantifies whether the observed prevalence of the clade is likely the result of chance. While ClaaTU can be applied to any phylogeny, the following investigations analyzed OTU trees to reduce tree complexity and subsequently increase statistical power.

We used ClaaTU to explore whether there are gut bacteria that are conserved across mammals. We analyzed fecal 16S rRNA sequences that were previously generated from 38 individuals spanning 32 different mammalian species and 10 orders (7). While these data represent a limited number of individuals, they provide a broad sampling of mammalian phylogenetic diversity. First, we determined that the evolutionary history of mammals is associated with the diversity of bacterial clades that comprise their gut microbiome (Supplemental File 1). We observed a strong and significant association between host Order, which served as a proxy for their evolutionary history, and the abundance - weighted (r^2^ = 0.44, p = 1e - 3) or presence - absence (r^2^=0.57, p=1e-3) Bray - Curtis dissimilarity among gut bacterial communities. Host feeding strategy (carnivory, omnivory, herbivory) more weakly associated with these measures of beta-diversity (r^2^ = 0.26, p = 1e - 3), although in the case of abundance - weighted beta - diversity, the strength of the association appears to be reduced by a subset of omnivores that group more closely with herbivores. Our results indicate that the evolutionary history of the host largely determines which clades are found in the mammalian gut, while the feeding strategy may determine which clades will predominate the community. These results are consistent with the patterns of phylosymbiosis observed elsewhere (6 – 8), but also underscore the importance of diet as a determining factor of which taxa dominate the gut microbiome (8, 22).

We then identified clades of bacteria that are conserved across mammals. Of the 8,086 clades harbored by the 38 individuals, 15 were ubiquitous among samples. However, the false discovery rate corrected p - values for these associations were insignificant because these clades appear near the root of the bacterial phylogeny and are thus likely ubiquitious by chance. Future work that considers the prevalence of these clades within a larger framework that includes non - mammalian lineages may reveal that they are indeed conserved within the mammalian gut. Similarly, deeper sequencing and expanded sampling per mammalian lineage may reveal the existence of universal and conserved clades in the mammalian gut. That said, we identified 38 more recently diverged clades that are found in a larger number of mammals than expected by chance (q < 0.2,). These conserved clades include members of the class Alphaproteobacteria, order Bacteroidales, the family Ruminococcaceae, and *Prevotella*. The observation that a clade within *Prevotella* is conserved among mammals (q = 0.04) is noteworthy because members of *Prevotella* produce short - chain fatty acids that contribute to intestinal health by serving as an energy source for host tissue, regulating inflammation, and promoting motility and blood flow (23). Additionally, Westernization among humans has been associated with a reduction of *Prevotella* (24).

To resolve deeper insight into how the gut microbiome relates to mammalian diversification, we next identified clades of bacteria that are conserved among distinct orders of mammals (Figure 2). Such clades may represent bacteria that are important to the unique physiological aspects of these sets of hosts. We constrained this analysis to the Carnivora, Artiodactyla, and Primates, as they were the only orders for which more than three host species were sampled. We identified 322, 591, and 633 clades that were conserved (q < 0.2) in these orders, respectively. Of these, 107, 255, and 245 were also uniquely present in their respective order. For example, all primates included in our study contained a clade of *Prevotella* (q = 0.0064) that was also unique to Primates, and consequently distinct from the *Prevotella* clade conserved across mammals. Similarly, a clade within *Faecalibacterium prausnitzii* is exclusive to and conserved among primates (45% of lineages) included in this study (q = 5.7e - 4), which is noteworthy given that *F. prausnitzii* is an abundant, butyrate - producing member of the healthy human gut microbiome whose underrepresentation in the gut is linked to gastrointestinal disorders (25, 26). Furthermore, a clade within *Turicibacter* was conserved in both Artiodactyla (66% of lineages) and Carnivora (71% of lineages), and missing from all other hosts. This clade may interact with mammalian traits that evolved in the ancestor of Artiodactyla and Carnivora and were potentially lost in Black Rhino, Zebras, and Horse. In support of this hypothesis, prior work in mice identified genetic loci that strongly associate with *Turicibacter* (27).

**Figure 2.**
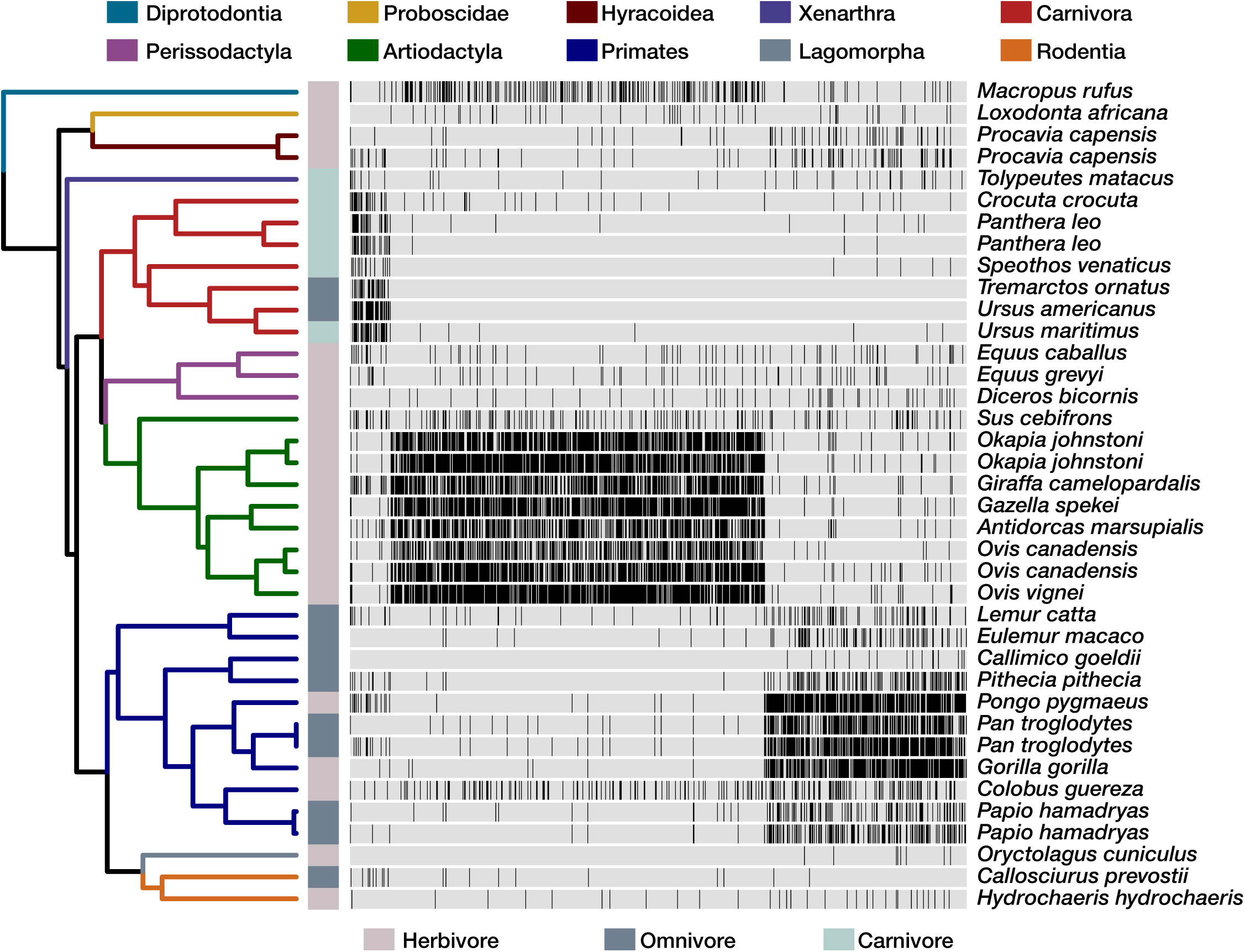
The phylogenetic distribution of conserved bacterial clades reveals associations between gut microbiota and mammalian evolutionary history. The 865 clades that are conserved in at least one mammalian Order (q < 0.2) and are not associated with dietary strategy are plotted as columns in a heatmap that illustrates their occurrence across mammalian lineages as black ticks. This includes 38 clades that are conserved across the mammals considered in this study. The dendrogram illustrates the evolutionary relationship among mammals, where edges are colored by order and dietary strategy is indicated adjacent to the tips.

We similarly identified clades that were conserved among mammals that were grouped by their dietary strategy to verify that the aforementioned patterns of clade conservation were not due to the potential confounding factor of host dietary strategy (Supplemental File 2). In doing so, we corroborated prior work by finding that omnivores carry clades that appear to be specialists to either herbivorous or carnivorous diets (8). However, unlike prior work, we also identified conserved clades of gut bacteria that are unique to omnivores, indicating that omnivore - specialist bacteria may exist.

### Conserved gut microbiota exhibit evolutionary patterns consistent with selection

We used several phylogenetic methods to discern whether natural selection could be influencing the conservation of bacterial clades across mammals. First, we assessed whether conserved clades are clustered across the bacterial phylogeny, which would indicate that the traits that result in a clade‘s conservation could arise through adaptive radiations or exaptation, be subject to environmental filtering, or improve dispersal (28). We calculated the phylogenetic distance between all pairs of conserved clades and used a Kolmogorov - Smirnov test to determine if the distribution of these distances differs from a bootstrapped distribution of clades randomly sampled from across the same phylogeny. Our analysis uncovers support for the phylogenetic clustering of clades conserved across either Carnivora (p = 4.5e - 10) or Primates (p = 0.03), but not Artiodactyla. In addition, considering only those clades that are both conserved and unique to Primates (p = 0.015), Artiodactyla (p = 0.015), or Carnivora (p = 2.2e - 9) reveals evidence of phylogenetic clustering. Furthermore, we tested whether clades that are conserved across dietary strategies are clustered, finding that carnivore (p = 6.7e - 7) and herbivore (p = 1.7e - 7) conserved clades are clustered, while omnivore conserved clades are not (p = 0.11). However, support for clustering improved when considering only those clades that are both conserved and unique to each of the dietary strategies (carnivore p = 2.6e - 10; herbivore p = 6.7e - 7; omnivore p = 2.2e - 16). We determined that signal propagation of clade conservation among closely related clades did not substantially affect these results by reconducting the analysis after excluding conserved clades whose parents were also conserved. Our finding of phylogenetic clusters of clade conservation indicates that some bacterial lineages are more likely to become conserved than others, possibly because their ancestors evolved a genetic background that potentiated the emergence of traits that improved the fitness of these lineages or resulted in their selection by the host.

We next reasoned that host filtering of bacterial lineages could give rise to independent clades that are exclusively conserved in distinct mammalian orders. An emerging body of research indicates that mammals and at least a limited number of their gut microbiota manifest patterns of co - diversification (8, 12). We hypothesize that some gut microbiota may be subject to a related process, wherein mammalian evolution is associated with the ecological sorting of bacterial lineages that derived from the same ancestor, but which innovated distinct traits that were then subject to selection, phylogenetic redundancy, and conservation among distinct groups of mammals. To explore the existence of such processes, we used parafit (29) to identify 1,171 clades of bacteria that manifest patterns of codiversification with their mammalian hosts (q < 0.05). Of these, 31 clades were conserved in at least one mammalian Order, indicating that some clades may become conserved within a group of mammals and subsequently are subject to coevolutionary processes. This includes clades within Clostridiales, which prior work found to be codiversifying with mammals (8), as well as a clade within *Prevotella*. We also identify evidence for codiversification of a clade within BS11, which prior work found to be cosmopolitan to ruminants (30), as well as a clade within Burkholderiales, which is known to contain lineages that metabolize toxic dietary xenobiotics, such as oxalate (31) (Supplemental File 3). Moreover, we identify 248 clades that codiversify with mammals and were not themselves conserved among any mammalian order, but gave rise to multiple descendant clades that were exclusive to and conserved within distinct orders. For example, we identify a clade within the Bacteroidales that gave rise to a clade conserved among the Artiodactyla as well as a distinct clade that is conserved among the Primates and annotated as being a member of *Prevotella* (Figure 3). These results support the hypotheses that at least some anciently integrated members of the mammalian microbiome may have diversified in concert with mammalian evolution, and that distinct sets of their descendants can become independently conserved in discrete groups of mammals. Collectively, these findings indicate that the evolution of at least some gut microbiota is linked to the evolution of their mammalian hosts.

**Figure 3.**
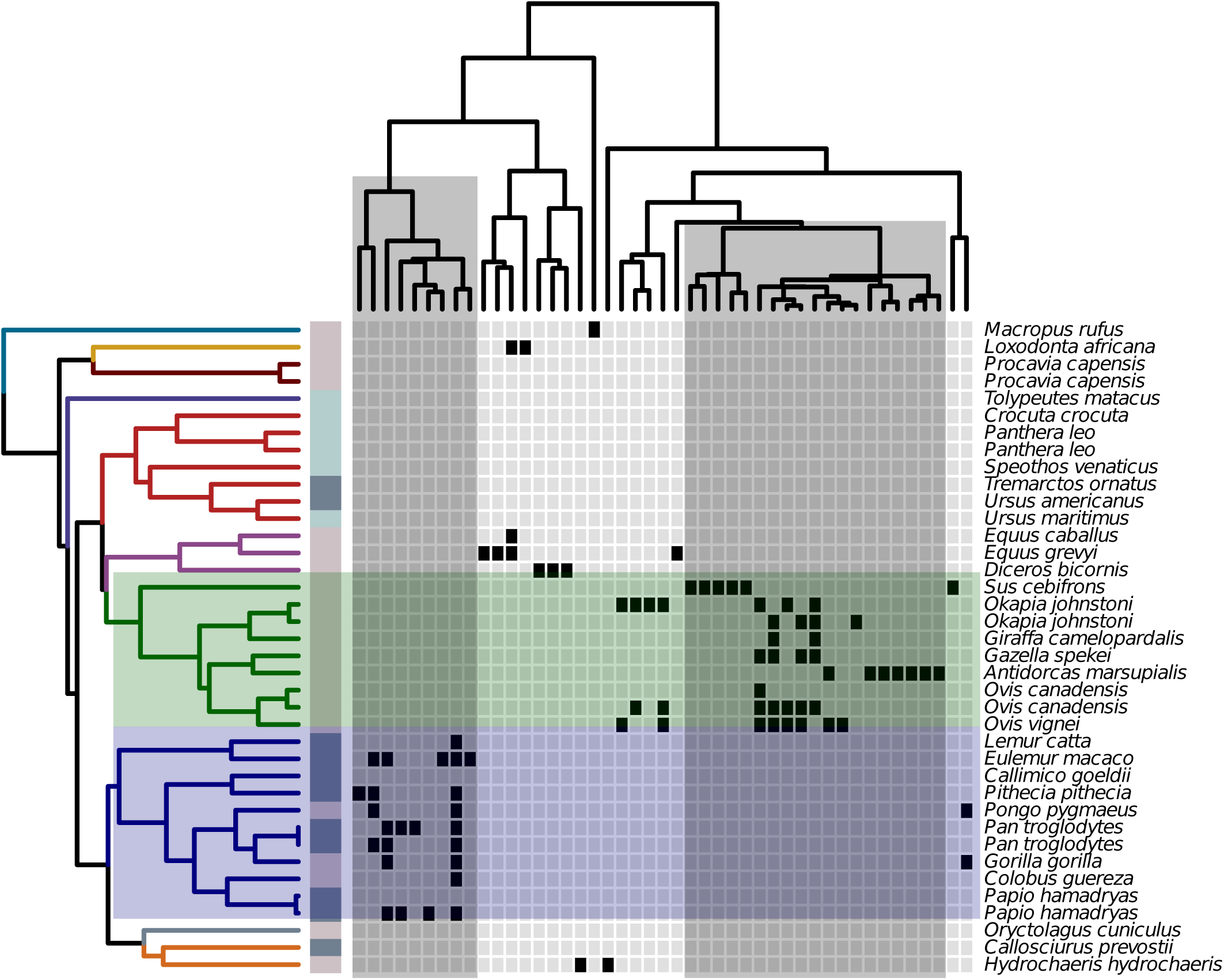
A co - diversifying clade within the Bacteroidales contains subtending clades that are unique to and conserved among discrete mammalian Orders. The evolutionary history of the OTUs in this co - diversifying (parafit; q < 0.05) bacterial clade is illustrated through the upper cladogram, while the left - hand cladogram relates mammalian lineages as in Figure 2. Black cells in the heat map indicate that the OTU was detected in a particular individual. Two subclades, highlighted in grey, are conserved among and unique to either the Artiodactyla (green) or Primates (blue).

### Lineage - specific variation of conserved clades in hominids

To clarify how the gut microbiome has diversified in association with human evolution (32), we applied ClaaTU to a deeply sequenced set of gut microbiome samples collected from a large number of hominid individuals. Following (33), we combined two large 16S fecal microbiome datasets that were prepared using matched molecular methods: one consisting of wild chimpanzees (n = 146), gorillas (n = 177), and bonobos (n = 69), and another consisting of humans from the United States of America (n = 314), Venezuela (n = 100), and Malawi (n = 114) (34). We assembled these data into a bacterial phylogeny with 69,517 clades. Due to the biological replication per lineage, these data afford accurate resolution into lineage - specific features of the microbiome.

The gut microbiome clade diversity among primates is associated with their evolutionary history (Figure 4) (6). For example, the clade Bray - Curtis dissimilarity stratified individuals based on their host species (adonis; R^2^ = 0.39, p < 0.001) as well as whether they were either non - human (chimp, bonobo, gorilla) or human (R^2^ = 0.28, p < 0.001). Furthermore, the clade Bray - Curtis dissimilarity correlated with the phylogenetic distance spanning samples (mantel test; R^2^ = 0.86, p < 1e - 4). Clade alpha - diversity also varied significantly across species, with notably depreciated Shannon entropy in gorillas and western humans. Collectively, these results corroborate prior work (6) that uncovered evidence of phylosymbiosis among primates and their gut microbiota, despite the fact that our respective studies vary in terms of host species and individuals sampled in addition to different considerations of microbial taxonomy.

**Figure 4.**
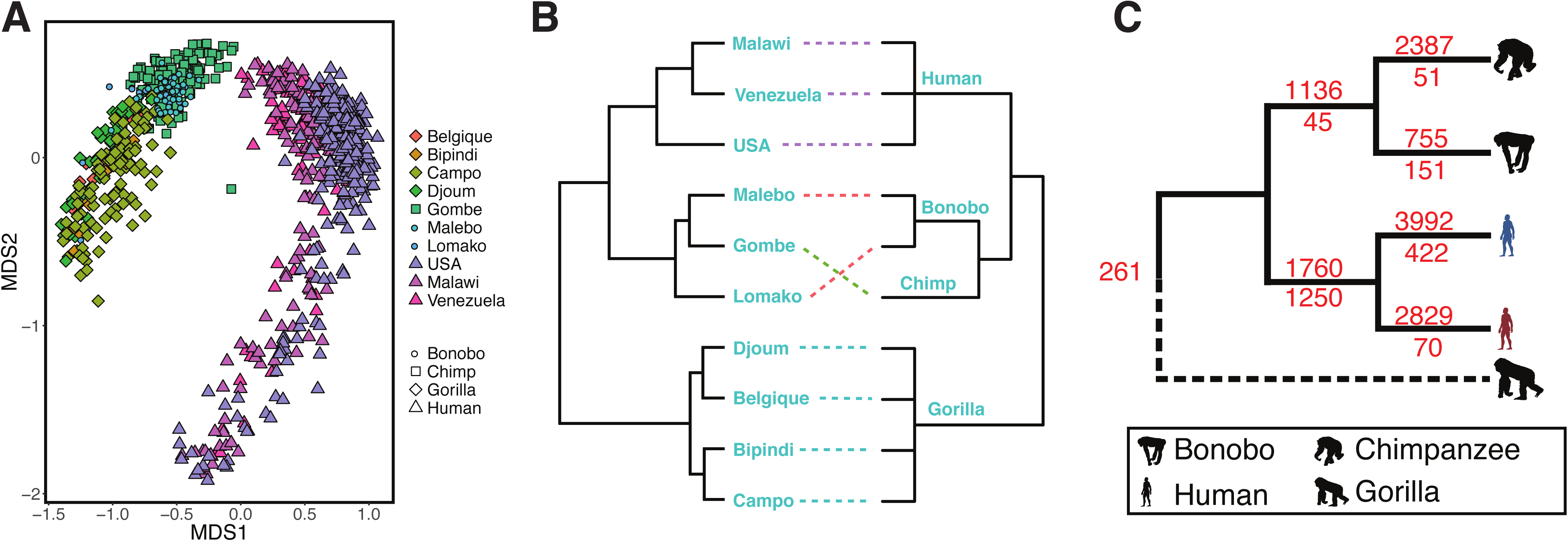
Primate gut microbiomes have diversified in a manner correlated with their evolutionary history. (A) A non - metric multi - dimensional scaling plot illustrates the significant differences in clade beta - diversity among primate groups (adonis; p < 0.001). (B) The dendrogram relating groups of primates by their microbiome clade beta - diversity (left) is significantly correlated with the phylogenetic distance spanning these same groups (right; mantel; p < 1e - 4). (C) A parsimony imputation of the acquisition (black numbers) and loss (red numbers) of conserved clades among primates that are grouped by their evolutionarily relationships finds that humans have a disproportionately low number of clades that are otherwise conserved among primates, and that this effect is amplified in Western humans (blue) as compared to Non-western Humans (red).

We also found that hominid species vary in terms of which bacterial clades are conserved in their microbiomes. We identified 18,942 clades that were conserved (fdr < 0.01) in at least one group of hominids (chimpanzee, gorilla, bonobo, western human, non - western human), only 261 of which were ubiquitously conserved (Supplemental File 4. An additional 1,250 were conserved among the wild apes, while 1,760 clades were conserved in western and non - western humans. Of these, 265 and 352 clades were exclusive (i.e., unobserved outside of the group) to non - human primates and humans respectively. For example, 66 clades associated with the short - chain fatty acid producing family Lachnospiraceae were exclusively conserved in humans and absent in the other primates (35). Non - human hominids contained 43 exclusively conserved clades associated with the polyphenol metabolizing family Coriobacteriacea (36). Western humans had the greatest number of conserved as well as exclusive and conserved clades. They also harbored the highest fraction of clades that were uniquely conserved (i.e., present in other lineages but not conserved; 55%). These results indicate that western humans share a substantial portion of their microbiome that is distinct from non - western humans and the non - human primates. This includes 508 conserved clades from the genus *Bacteroides*, 107 from *Ruminococcus*, and 54 from *Akkermansia*. These results are consistent with previous reports that indicate that diets rich in fats, such as the standard western diet, support microbiomes high in *Bacteroides* (37). Conversely, non - western individuals harbored a large number of uniquely conserved clades from the genera *Prevotella* (733 clades), *Streptococcus* (79 clades), *Lactobacillus* (57 clades), and *Bifidobacterium* (56 clades). Chimpanzees, bonobos, and gorillas also harbored several clades associated with the genus *Prevotella* (128, 80, 63 clades respectively), while western humans only contained three, indicating that substantial conserved clade diversity of *Prevotella* is a feature missing from western humans. Our results are consistent with previous observations that *Prevotella* abundance is associated with dietary fiber intake (37), and further suggest that consumption of low fiber diets, such as the standard western diet, may result in decreased diversity of *Prevotella* and increased diversity of *Bacteroides* clades in the gut. Collectively, this analysis indicates that changes in lifestyle, environment, or genetics that are associated with westernization have occurred in concert with changes to the suite of conserved bacteria that occupy the gut.

Given these patterns of clade conservation across hominids, we used a parsimony approach to identify gut bacterial clades that have become conserved (i.e., gains) or that are no longer conserved (i.e., losses) along specific hominid lineages. For this analysis, gorilla was used as an out - group, which prevented the assessment of gains and losses along this lineage. We observed a relatively extensive gain of conserved clades by each species during the course of hominid evolution (Figure 4). Gains in all lineages outpaced losses potentially due to additional niches opening in the gut during speciation, dietary transition, or altered habitat.

The human gut microbiome dramatically differs from other hominids in terms of conserved clades. For example, humans lost disproportionately large number of hominid conserved clades. Westernized humans exemplify this trend. These clades include members of common human gut genera, such as *Prevotella* (45 clades), *Methanobrevibacter* (19 clades), and *Bifidobacterium* (4 clades) (Supplemental File 5). Humans also gained conserved clades associated with the genera *Bacteroides* (64 clades), *Bifidobacterium* (32 clades), and *Ruminococcus* (26 clades). These results agree with prior research that indicated that the abundance of *Ruminococcus, Bifidobacterium,* and *Bacteroides* differentiated non - human from human primates (33). Our findings suggest that humans evolved in conjunction with substantial alterations to the factors that impact gut microbiome clade conservation. While the underlying processes driving these patterns are not known, they potentially include: 1) interlineage variation in ecology and environment, including differences in shelter and sanitation that could affect microbial metacommunity exposure (38); 2) genomic evolution, as genotype - microbiome interactions have been described in humans (15); 3) aspects of behavior or diet that may influence microbial dispersal and growth in the gut; and 4) cryptic study effects that bias the resolution of microbial lineages in specific host groups. An expanded sampling of primate lineages, across diverse populations, coupled with rich metadata would help determine whether these processes contribute to clade conservation in primates.

## Discussion

Phylogenetics empowers the characterization of microbial community diversity (39 – 42). Here, we used phylogeny to resolve microbial taxonomic units that are based on shared ancestry and shared ecology. Our approach identified monophyletic clades of microbial taxa that were unexpectedly prevalent across mammals. These conserved clades of gut microbiota are important to identify because they can potentially reveal the processes that influence the ecological and evolutionary diversification of the mammalian gut microbiome. For example, selective factors may influence taxa that are conserved across communities, as their distributions are unlikely to occur through random processes (16). It is unclear if these enteric bacterial clades are conserved because they are either gut generalists that are effective at dispersal, or alternatively because their ancestor integrated into the mammalian gut microbiome, elicited a beneficial effect for the host, and was subsequently retained as mammals diversified, possibly due to natural selection. Since many of these animals were kept in captivity and potentially in close proximity, it is possible that these clades are transient and environmentally acquired. Future work should discern the mechanisms underlying their high prevalence among mammals.

Relatively few clades were present in all mammalian lineages, and most of the conserved clades identified were not fully present across all individuals within a considered group. Discrete lineages of gut bacteria have convergently evolved ecologically relevant traits, such as glycosyl hydrolases (43), which could account for an absence of fully present distributions of conserved clades across hosts; if selection is acting on these clades, it is likely doing to at the level of these traits. Alternatively, mammalian evolution may have yielded changes in the selective regime for specific microbiome traits. Such changes would reduce the ubiquity of these clades across mammals. Future work should consider expanding the numbers of individuals sampled from each host lineage to improve the resolution of species - specific associations with the microbiome, especially given that enterotypic variation (44) or low depth of sampling could create the signature of a clade‘s absence when few individuals are interrogated within a host species. Indeed, in our expanded analysis of the primates, we resolved the complete absence of some clades in humans that are otherwise conserved across hominids despite an extensive sampling of individuals. These findings support the hypothesis that there exist host lineage - specific dependencies on the gut microbiome, but further work that controls for facility, geographic, and dietary effects is needed.

Our analysis only considers bacteria from stool and does not consider the environmental metacommunity from which animals sample their microbiomes (11). This matters for several reasons. First, this metacommunity may have changed over the course of host evolution. Thus, our findings do not necessarily indicate that the microbiome has co - evolved with their hosts, or even that the observations made here were necessarily present throughout the evolution of the various lineages being studied. For example, it is possible that the distributions of taxa observed across these hosts have been influenced by contemporary shifts in the metacommunity population, such that the microbes that associated with host ancestors were distinct from those that associate with extant taxa. However, given the phylogenetic patterns in our data, we may expect that the traits that make contemporary microbes successful at colonizing a large number of hosts were inherited from those bacteria that successfully colonized ancestral hosts. Second, it is possible that metacommunity variation contributes to the differences observed between taxa. For example, perhaps the clades that stratify humans from non - human primates are not ubiquitous to the metacommunities of these two sets of hosts. Consequently, the conserved clades that stratify hosts may do so because they possess traits that 1) enable their success in their respective metacommunity and 2) enable their frequent migration into the guts of their respective hosts. Third, while there appears to be some evidence of inheritance in our results, we note that this does not necessarily indicate that direct, vertical transmission of microbiota has occurred between generations. Indeed, patterns of microbiome inheritance may result from vertical transmission of host genotype that selects for specific metacommunity assemblages.

Our finding of conserved clades is consistent with recent efforts to ascribe periods of microbial evolution to the emergence of specific traits (19, 42, 45). Indeed, phylogenetic redundancy results when different phylogenetic lineages can be interchanged in a community because they execute similar ecological roles (16). Our informatic procedure identifies monophyletic clades that contain redundant lineages, indicating that the origin of the trait in question likely emerged in or prior to the clade‘s ancestor. However, limitations in the phylogenetic breadth of the data analyzed here may result in inaccurate imputation of a trait‘s evolutionary origin. Future work should seek to map traits onto comprehensive phylogenies to assess when traits arose and which specific traits are likely contributing to clade conservation. This will consequently facilitate empirical investigations of the role of these traits in the operation of the gut microbiome, microbial dispersal, and host fitness.

## Materials and Methods

### Identification of Ecophylogenetic Taxonomic Units

We developed ClaaTU (https://github.com/chrisgaulke/Claatu), an algorithm that quantifies the abundance of monophyletic clades of taxa across a set of communities and optionally identifies clades that are more prevalent than expected by chance. ClaaTU conducts a brute - force algorithm root - to - tip traversal of a phylogenetic tree and quantifies the abundance of each monophyletic clade in each community by summing the abundances of subtending lineages. Consequently, ClaaTU resolves the ecological distribution of monophyletic groups of taxa. To determine if a clade is conserved, ClaaTU converts abundances to presence - absence data and then quantifies each clade‘s prevalence across a set of samples. To ascertain if the observed prevalence is greater than that expected by chance, ClaaTU conducts a phylogenetic permutation test (46). Specifically, the observed phylogenetic tip – to - community labels are randomly shuffled, such that the underlying prevalence distribution remains fixed, while the associated lineages are altered. This random permutation of the data occurs multiple times (1000 for the mammal - wide study and 100 for the primate study due to limitations of tree size) to produce a bootstrapped prevalence distribution for each clade. A z - test determines if the clade‘s observed prevalence is significantly greater than the bootstrapped null distribution. ClaaTU assigns a taxonomic label to each clade by identifying the most granular taxonomic assignment shared by all subtending lineages.

#### Analysis of Mammalian Microbiome Samples

We analyzed publicly available data. First, data from thirty-one animals representing ten taxonomic orders (7) was downloaded from MG - RAST (mgp 113, mgp 114). Reads were clustered into operational taxonomic units (OTUs) using pick_open_reference_otus.py in QIIME (47) with UCLUST (48) using a 97% identity threshold against the greengenes database (13_8). Taxonomy was inferred for each OTU using assign_taxonomy.py in QIIME with default parameters.

Second, a dataset consisting of 146 wild chimpanzees, 69 bonobos, and 177 gorillas from several field sites (33) was downloaded from http://web.biosci.utexas.edu/ochman/moeller_data.html and filtered to trim low quality bases (q < 25) using split_libraries.py in QIIME. A separate dataset (34) consisting of 528 humans - of which 314 were from the United States (western), and 114 and 100 were from Malawi and Venezuela, respectively (non - western) - was downloaded from MG - RAST (mgp401) The non - human and human datasets were combined, trimmed to a uniform sequence length of 99 bp, and collectively processed. OTU clustering and taxonomic annotation occurred as above. The resulting dataset was filtered to remove low frequency (present in less than nine samples) and low abundance (total abundance < 10 counts) OTUs.

#### Tree construction and clade diversity quantification

The QIIME - assigned OTU representative sequences were used to assemble phylogenetic trees for each data set. In the case of the primate data, these sequences were combined with the greengenes 97% identity set of full - length reference sequences to improve the phylogenetic accuracy of short sequence data, following (49, 50). Infernal (51) aligned sequences as in (49, 50). Alignment columns containing 50% gap characters were removed using filter_alignment.py in QIIME. FastTree constructed phylogenies using generalized time - reversible model (52). Phylogenies were pruned of greengenes reference sequences, subject to midpoint rooting, and processing by ClaaTU.

#### Statistical analyses

Statistical analyses were conducted using R. Bray - Curtis dissimilarity was calculated using vegan::vegdist, while vegan::diversity quantified Shannon entropy. Permutational Multivariate Analysis of Variance quantified the association between beta - diversity and categorical sample covariates (vegan::adonis; 1000 permutations). To determine if patterns of microbiome diversity were similar to the evolutionary history of primates, the Bray - Curtis disimilarity matrix was correlated to the phylogenetic distance matrix spanning all pairs of samples using the mantel function with 1,000 permutations. The hominid phylogeny was obtained from the 10kTrees (version 3) website (http://10ktrees.fas.harvard.edu/) (53). For all analyses, false discovery rate procedures corrected for multiple tests. All analytical software and results can be found at http://files.cgrb.oregonstate.edu/Sharpton_Lab/Papers/Gaulke_ISME_2017/.

## Acknowledgements

We thank Katherine Pollard, Jonathan Eisen, and Joshua Ladau for helpful discussions. The National Science Foundation (Grant 1557192) and institutional funds to TJS supported this work.

## Conflicts of Interest

TJS is a scientific advisor and holds equity in Resilient Biotics, LLC.

Supplementary information is available at ISMEJ‘s website

## References

1. Bäckhed F, G K, JJ F (2012) Host responses to the human microbiome. Nutr Rev 70(2):S14–S17.

2. Claus SP, Guillou H, Ellero-Simatos S (2016) The gut microbiota: a major player in the toxicity of environmental pollutants? npj Biofilms Microbiomes 2:16003.

3. Hooper L V, Littman DR, Macpherson AJ (2012) Interactions between the microbiota and the immune system. Science 336(6086):1268–73.

4. Cho I, Blaser MJ (2012) The human microbiome: at the interface of health and disease. Nat Rev Genet 13(4):260–70.

5. Cryan JF, Dinan TG (2012) Mind-altering microorganisms: the impact of the gut microbiota on brain and behaviour. Nat Rev Neurosci 13(10):701–12.

6. Brooks AW, et al. (2016) Phylosymbiosis: Relationships and Functional Effects of Microbial Communities across Host Evolutionary History. PLOS Biol 14(11):e2000225.

7. Ley RE, et al. (2008) Evolution of mammals and their gut microbes. Science 320(5883):1647–51.

8. Groussin M, et al. (2017) Unraveling the processes shaping mammalian gut microbiomes over evolutionary time. Nat Commun 8:14319.

9. Muegge BD, et al. (2011) Diet drives convergence in gut microbiome functions across mammalian phylogeny and within humans. Science 332(6032):970–4.

10. Clayton JB, et al. (2016) Captivity humanizes the primate microbiome. Proc Natl Acad Sci:201521835.

11. O’Dwyer JP, Kembel SW, Green JL (2012) Phylogenetic diversity theory sheds light on the structure of microbial communities. PLoS Comput Biol 8(12):e1002832.

12. Moeller AH, et al. (2016) Cospeciation of gut microbiota with hominids. Science (80-) 353(6297). Available at: http://science.sciencemag.org/content/353/6297/380 [Accessed May 7, 2017].

13. Goodrich JK, et al. (2014) Human Genetics Shape the Gut Microbiome. Cell 159(4):789–799.

14. Beaumont M, et al. Heritable components of the human fecal microbiome are associated with visceral fat. Genome Biol 17(1):189.

15. Blekhman R, et al. (2015) Host genetic variation impacts microbiome composition across human body sites. Genome Biol 16(1):191.

16. Shade A, Handelsman J (2012) Beyond the Venn diagram: the hunt for a core microbiome. Environ Microbiol 14(1):4–12.

17. Martiny AC, Treseder K, Pusch G (2013) Phylogenetic conservatism of functional traits in microorganisms. ISME J 7(4):830–8.

18. Philippot L, et al. (2010) The ecological coherence of high bacterial taxonomic ranks. Nat Rev Microbiol 8(7):523–9.

19. Barberán A, Caceres Velazquez H, Jones S, Fierer N (2017) Hiding in Plain Sight: Mining Bacterial Species Records for Phenotypic Trait Information. mSphere 2(4). Available at: http://msphere.asm.org/content/2/4/e00237-17 [Accessed August 24, 2017].

20. Mouquet N, et al. (2012) Ecophylogenetics: advances and perspectives. Biol Rev Camb Philos Soc 87(4):769–85.

21. Cohan FM, Kane M (2001) Bacterial Species and Speciation. Syst Biol 50(4):513–524.

22. Muegge BD, et al. (2011) Diet Drives Convergence in Gut Microbiome Functions Across Mammalian Phylogeny and Within Humans. Science (80-) 332(6032). Available at: http://science.sciencemag.org/content/332/6032/970 [Accessed May 7, 2017].

23. Flint HJ, Bayer EA, Rincon MT, Lamed R, White BA (2008) Polysaccharide utilization by gut bacteria: potential for new insights from genomic analysis. Nat Rev Microbiol 6(2):121–131.

24. De Filippo C, et al. (2010) Impact of diet in shaping gut microbiota revealed by a comparative study in children from Europe and rural Africa. Proc Natl Acad Sci 107(33):14691–14696.

25. Heinken A, et al. (2014) Functional metabolic map of Faecalibacterium prausnitzii, a beneficial human gut microbe. J Bacteriol 196(18):3289–302.

26. Miquel S, et al. (2013) Faecalibacterium prausnitzii and human intestinal health. Curr Opin Microbiol 16(3):255–261.

27. Theriot CM, Bowman AA, Young VB (2016) Antibiotic-Induced Alterations of the Gut Microbiota Alter Secondary Bile Acid Production and Allow for Clostridium difficile Spore Germination and Outgrowth in the Large Intestine. mSphere 1(1):e00045–15.

28. Horner-Devine MC, Bohannan BJM (2006) PHYLOGENETIC CLUSTERING AND OVERDISPERSION IN BACTERIAL COMMUNITIES. Ecology 87(sp7):S100–S108.

29. Legendre P, Desdevises Y, Bazin E (2002) A statistical test for host-parasite coevolution. Syst Biol 51(2):217–34.

30. Solden LM, et al. (2017) New roles in hemicellulosic sugar fermentation for the uncultivated Bacteroidetes family BS11. ISME J 11(3):691–703.

31. Stewart CS, Duncan SH, Cave DR (2004) Oxalobacter formigenes and its role in oxalate metabolism in the human gut. FEMS Microbiol Lett 230(1):1–7.

32. Schnorr SL, Sankaranarayanan K, Lewis CM, Warinner C (2016) Insights into human evolution from ancient and contemporary microbiome studies. Curr Opin Genet Dev 41. doi:10.1016/j.gde.2016.07.003.

33. Moeller AH, et al. (2014) Rapid changes in the gut microbiome during human evolution. Proc Natl Acad Sci 111(46):16431–5.

34. Yatsunenko T, et al. (2012) Human gut microbiome viewed across age and geography. Nature 486(7402):222–7.

35. Meehan CJ, Beiko RG (2014) A phylogenomic view of ecological specialization in the Lachnospiraceae, a family of digestive tract-associated bacteria. Genome Biol Evol 6(3):703–13.

36. Clavel T, Lepage P, Charrier C (2014) The Family Coriobacteriaceae. The Prokaryotes (Springer Berlin Heidelberg, Berlin, Heidelberg), pp 201–238.

37. David LA, et al. (2014) Diet rapidly and reproducibly alters the human gut microbiome. Nature 505(7484):559–63.

38. Ruiz-Calderon JF, et al. (2016) Walls talk: Microbial biogeography of homes spanning urbanization. Sci Adv 2(2). Available at: http://advances.sciencemag.org/content/2/2/e1501061 [Accessed May 29, 2017].

39. Lozupone C, Lladser ME, Knights D, Stombaugh J, Knight R (2011) UniFrac: an effective distance metric for microbial community comparison. ISME J 5(2):169–72.

40. Matsen FA, Evans SN (2013) Edge principal components and squash clustering: using the special structure of phylogenetic placement data for sample comparison. PLoS One 8(3):e56859.

41. Amend AS, et al. (2015) Microbial response to simulated global change is phylogenetically conserved and linked with functional potential. ISME J. doi:10.1038/ismej.2015.96.

42. Washburne AD, et al. (2017) Phylogenetic factorization of compositional data yields lineage-level associations in microbiome datasets. PeerJ 5:e2969.

43. Lozupone CA, et al. (2008) The convergence of carbohydrate active gene repertoires in human gut microbes. Proc Natl Acad Sci 105(39):15076–15081.

44. Moeller AH, et al. (2012) Chimpanzees and humans harbour compositionally similar gut enterotypes. Nat Commun 3:1179.

45. Martiny JBH, Jones SE, Lennon JT, Martiny AC (2015) Microbiomes in light of traits: A phylogenetic perspective. Science (80-) 350(6261). Available at: http://science.sciencemag.org/content/350/6261/aac9323 [Accessed May 7, 2017].

46. Martin AP (2002) Phylogenetic approaches for describing and comparing the diversity of microbial communities. Appl Environ Microbiol 68(8):3673–82.

47. Caporaso JG, et al. (2010) QIIME allows analysis of high-throughput community sequencing data. Nat Methods 7(5):335–6.

48. Edgar RC (2010) Search and clustering orders of magnitude faster than BLAST. Bioinformatics 26(19):2460–1.

49. Sharpton TJ, et al. (2011) PhylOTU: a high-throughput procedure quantifies microbial community diversity and resolves novel taxa from metagenomic data. PLoS Comput Biol 7(1):e1001061.

50. O’Dwyer JP, Kembel SW, Sharpton TJ (2015) Backbones of evolutionary history test biodiversity theory for microbes. Proc Natl Acad Sci 112(27):201419341.

51. Nawrocki EP, Eddy SR (2013) Infernal 1.1: 100–fold faster RNA homology searches. Bioinformatics 29(22):2933–2935.

52. Price MN, Dehal PS, Arkin AP (2010) FastTree 2--approximately maximum-likelihood trees for large alignments. PLoS One 5(3):e9490.

53. Arnold C, Matthews LJ, Nunn CL (2010) The 10kTrees website: A new online resource for primate phylogeny. Evol Anthropol Issues, News, Rev 19(3):114–118.

54. Kohl KD, Weiss RB, Cox J, Dale C, Dearing MD (2014) Gut microbes of mammalian herbivores facilitate intake of plant toxins. Ecol Lett 17(10):1238–46.

